## Additional Data 1 for "The stringent stress response controls proteases and global regulators under optimal growth conditions in *Pseudomonas aeruginosa*"

**Appendix 1. *P. aeruginosa* ppGpp synthase RelA was under dual-promoter control**

Based on promoter prediction (1) and transcriptional profiling (2), the ppGpp synthase *relA* is in an operon with the cysteine synthase *cysM* and RNA methyltransferase *ygcA* (*SI Appendix;* Fig S1A). Intriguingly, although these formed an operon, genome-wide transcription start site mapping revealed two transcriptional start sites and several σ factor binding sites including RpoH (σ^32^) and RpoS (σ^38^) upstream of both *relA* and *cysM* (2-4). To further investigate each promoter activity, the upstream promoter region of *relA* and *cysM* was transcriptionally fused to the fluorescent reporter gene *mCherry*. Since we previously showed that the promoter activity of *relA* is very low when chromosomally complemented (as evidenced by non-lethal expression of *relA* in a *relA/spoT* mutant - resembling a *spoT* knockout) (5), we created the fusion constructs on a plasmid. While fluorescence levels were below a reliable signal strength during exponential growth for the *relA* and *cysM* promoter, *relA* promoter activity significantly increased in the wild type when cells transitioned into stationary phase after 8 h (2.75-fold) (*SI Appendix;* Fig. S1C).

Since many amino acid biosynthesis pathways have been identified that respond to amino acid starvation and ppGpp induction (6), we further investigated promoter activity upon amino acid (i.e., arginine) starvation. Reducing the amount of arginine in the medium had no effect on wild-type growth, however, the significant signal for *relA* promoter activity was only detected after 12 h cf. to 8 h as mentioned above. The signal for *cysM* promoter activity was slightly affected and significant after 15 h in 6.5 μM cf. 14 in 12.5 μM arginine (*SI Appendix;* Fig S1B, S1C). This prompted us to further investigate promoter activity in an arginine auxotrophic PAO1 mutant (Δ*argB*). While the mutant was not able to grow without the supplementation of at least 1 μM arginine in minimal medium, 12.5 μM restored growth similar to wild type levels (*SI Appendix;* Fig S1B). In Δ*argB* the *relA* promoter activity significantly increased after 11 h (2.45-fold) in 12.5 μM arginine and also the *cysM* signal significantly increased at 14 h (2.60-fold). Intriguingly, the promoter activity of both promoters was significantly shifted at 6.25 μM arginine, being significant at 9 h (*relA*) and 11 h (*cysM*) (*SI Appendix;* Fig S1B, S1C). Together, these results suggested that both the *relA* and *cysM* promoter are involved in activation of the RelA synthase gene expression during amino acid starvation in *P. aeruginosa*. In addition, both promoters were required for maximum accumulation of RelA during stationary phase, as suggested by the increased signal intensity when both promoters were fused together (*SI Appendix;* Figure S1D, S1E), as also previously observed by Nakagawa *et al*. in *E. coli* (7). Critically although gene organization is quite different in *P. aeruginosa* and *E. coli*, with two genes between the two promoters in the former, the general principle of a distal, inducible promoter and a constitutive proximal promoter appears to have been conserved.

With regards to the regulation of *spoT*, which is in the same operon as the ω subunit *rpoZ* of the polymerase in *E. coli* (8), we found that based on the promoter prediction and the transcriptional start site mapping, *spoT* was also in an operon with *rpoZ* in *P. aeruginosa* PAO1. The small ω subunit aids in in the final step of the core polymerase assembly, interacts with ppGpp binding at the β’-ω interface, and allows for alteration of gene expression (9). The ω subunit is also required for RNA polymerase stability and transcriptional specificity (9, 10). To study its promoter activity, we fused the PAO1 *rpoZ-spoT* promoter to the fluorescent reporter gene *gfp* and measured fluorescence during growth in minimal medium. The fluorescence levels increased with increasing cell density and showed strong expression throughout the cell cycle (data not shown). The continuous expression of *spoT* throughout cell growth is consistent with its role in metabolizing ppGpp (to prevent excess buildup), but might also indicate that SpoT has other hitherto unknown important functions.
